# Distinct Nuclear and Cytoplasmic Transcriptomic Signatures Reveal Transcription Alone Is Insufficient to Determine Glucose-Induced Transcriptomic Dynamics

**DOI:** 10.1101/2024.12.22.629974

**Authors:** Atefeh Bagheri, Jinsil Kim, Peng Jiang

## Abstract

Glucose profoundly influences cellular transcriptomes, but whether these changes are primarily driven by transcription remains unclear. Traditional bulk RNA sequencing, which interrogates total mRNA from whole cells, obscures distinct dynamics of nuclear and cytoplasmic transcriptomes. Nuclear RNA levels primarily reflect transcriptional activity, whereas cytoplasmic RNA levels are shaped by both transcription and post-transcriptional processes, such as RNA export, stability, and degradation. In this study, we systematically investigate glucose-induced transcriptomic responses in a subcellular location- and cell type-specific manner using three cell lines: FHC (normal colonic epithelial cells), MCF10A (normal breast epithelial cells), and MCF7 (metastatic breast cancer cells). Our findings reveal that, although nuclear and cytoplasmic mRNA levels show strong global correlations, glucose-induced changes in mRNA abundance exhibit minimal concordance between the nucleus and cytoplasm. Additionally, glucose-induced changes in exon inclusion levels often diverge between the nucleus and cytoplasm, underscoring the importance of post-transcriptional processes in shaping the cytoplasmic transcriptome response to glucose level changes. Glucose-induced differentially expressed genes (DEGs) and differentially spliced exons (ΔPSI) are enriched in distinct pathways exhibiting unique enrichment patterns depending on the subcellular location and cell line. These findings underscore the complexity of glucose-induced transcriptomic regulation, demonstrating that transcription alone is insufficient to explain the observed dynamics.

## Introduction

Glucose is a key metabolic regulator that profoundly influences gene expression, impacting vital processes such as cell growth, proliferation, and survival (Zhao et al. 2008). While the transcriptional responses to glucose are well-documented (Meugnier et al. 2007), much less is understood about the role of post-transcriptional mechanisms—such as mRNA export, stability, and alternative splicing—in shaping the transcriptome. Traditional approaches to studying glucose-induced transcriptomic changes have largely relied on bulk RNA sequencing(Wang et al. 2009; Pang et al. 2016; Webert et al. 2022), which measures total mRNA levels without distinguishing between nuclear and cytoplasmic compartments. As mRNA must be exported from the nucleus to the cytoplasm for translation, nuclear mRNA abundance does not necessarily correlate with cytoplasmic mRNA levels. Moreover, post-transcriptional processes (Corbett 2018), including mRNA stability and alternative splicing, critically shape the functional mRNA pool in the cytoplasm.

Recent advances in subcellular location-specific transcriptomics have underscored the importance of separately analyzing nuclear and cytoplasmic transcriptomes to unravel the distinct regulatory mechanisms operating in each compartment (Zaghlool et al. 2021; Hurni et al. 2022). However, few studies have systematically examined how glucose influences transcriptomic dynamics in a compartment-specific manner or whether these effects vary across different cell types.

In this study, we address these gaps by investigating the nuclear and cytoplasmic transcriptomic responses to glucose in three distinct cell lines: FHC (normal colonic epithelial cells), MCF10A (normal breast epithelial cells), and MCF7 (metastatic breast cancer cells). Through subcellular location-specific RNA sequencing and pathway enrichment analysis, we aim to disentangle the contributions of transcriptional and post-transcriptional regulation to glucose-induced transcriptomic changes.

Our findings reveal that glucose-induced changes in mRNA abundance exhibit minimal concordance between the nuclear and cytoplasmic compartments, highlighting the critical role of post-transcriptional processes such as mRNA export and stability in shaping cytoplasmic transcriptomes. Furthermore, we demonstrate that glucose-induced alternative splicing changes are highly compartment- and cell type-specific, further emphasizing the complexity of transcriptomic regulation.

Pathway enrichment analysis reveals distinct biological pathways affected by glucose in a compartment- and cell type-specific manner. By dissecting the compartmental and cell type-specific transcriptomic responses to glucose, our study provides critical insights into the nuanced regulation of cellular metabolism and underscores the need for more targeted approaches in transcriptomic analysis.

## Results

### Discrepancy between nuclear and cytoplasmic mRNA abundance changes

Numerous studies have investigated how cells respond to changes in glucose levels, with most relying on bulk RNA sequencing, which typically measures mRNA levels in whole cells, encompassing both nuclear and cytoplasmic compartments. While mRNA is transcribed in the nucleus and subsequently exported to the cytoplasm for translation, it remains unclear whether changes in mRNA abundance in the nucleus directly correspond to changes in the cytoplasm. In this study we aim to explore the relationship between nuclear and cytoplasmic mRNA abundance changes and to determine whether these correlations are dependent on cell type. We analyzed three cell lines: FHC, MCF10A, and MCF7.

We first calculated Spearman’s rank correlations to assess the relationship between nuclear and cytoplasmic mRNA relative abundance across the three cell lines under two conditions: normal glucose (5 mmol/L) and high glucose (25 mmol/L). The analysis revealed a strong global correlation between nuclear and cytoplasmic mRNA levels, with Spearman’s correlation coefficient (Rho) exceeding 0.97 for all cell lines and conditions (**Figure 1**).

**Figure 1.**
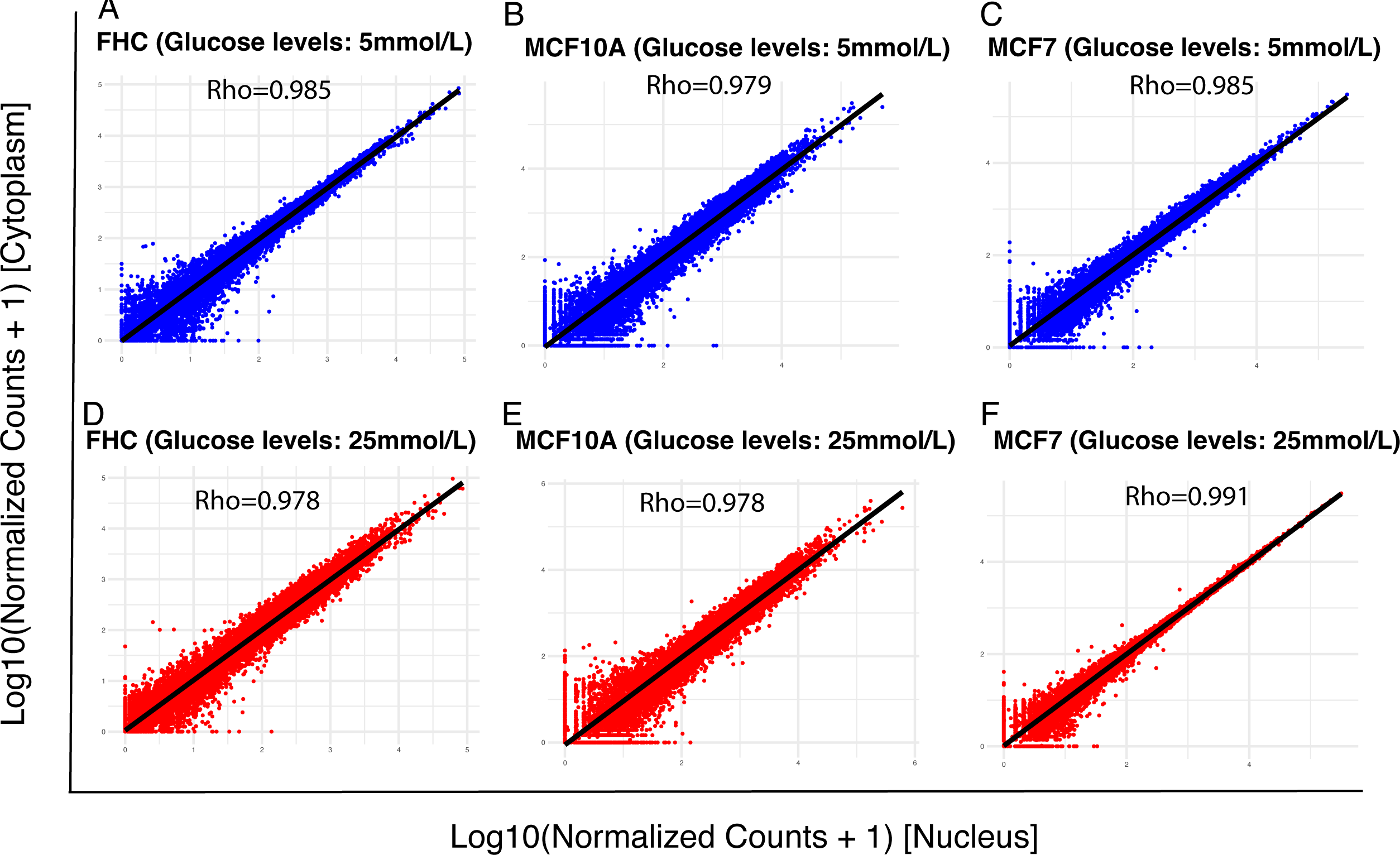
Spearman’s rank correlations of mRNA abundance between the nucleus and cytoplasm. (A) FHC cell line with low glucose level. (B) MCF10A cell line with low glucose level. (C) MCF7 cell line with low glucose level. (D) FHC cell line with high glucose level. (E) MCF10A cell line with high glucose level. (F) MCF7 cell line with high glucose level. The Spearman’s correlation coefficient (Rho) is indicated in each figure.

Next, we investigated whether transcriptomic changes in the cytoplasm due to varying glucose levels reflected those observed in the nucleus. Surprisingly, our results showed poor correlation between glucose-induced changes in nuclear and cytoplasmic mRNA abundance (**Figure 2**).

**Figure 2.**
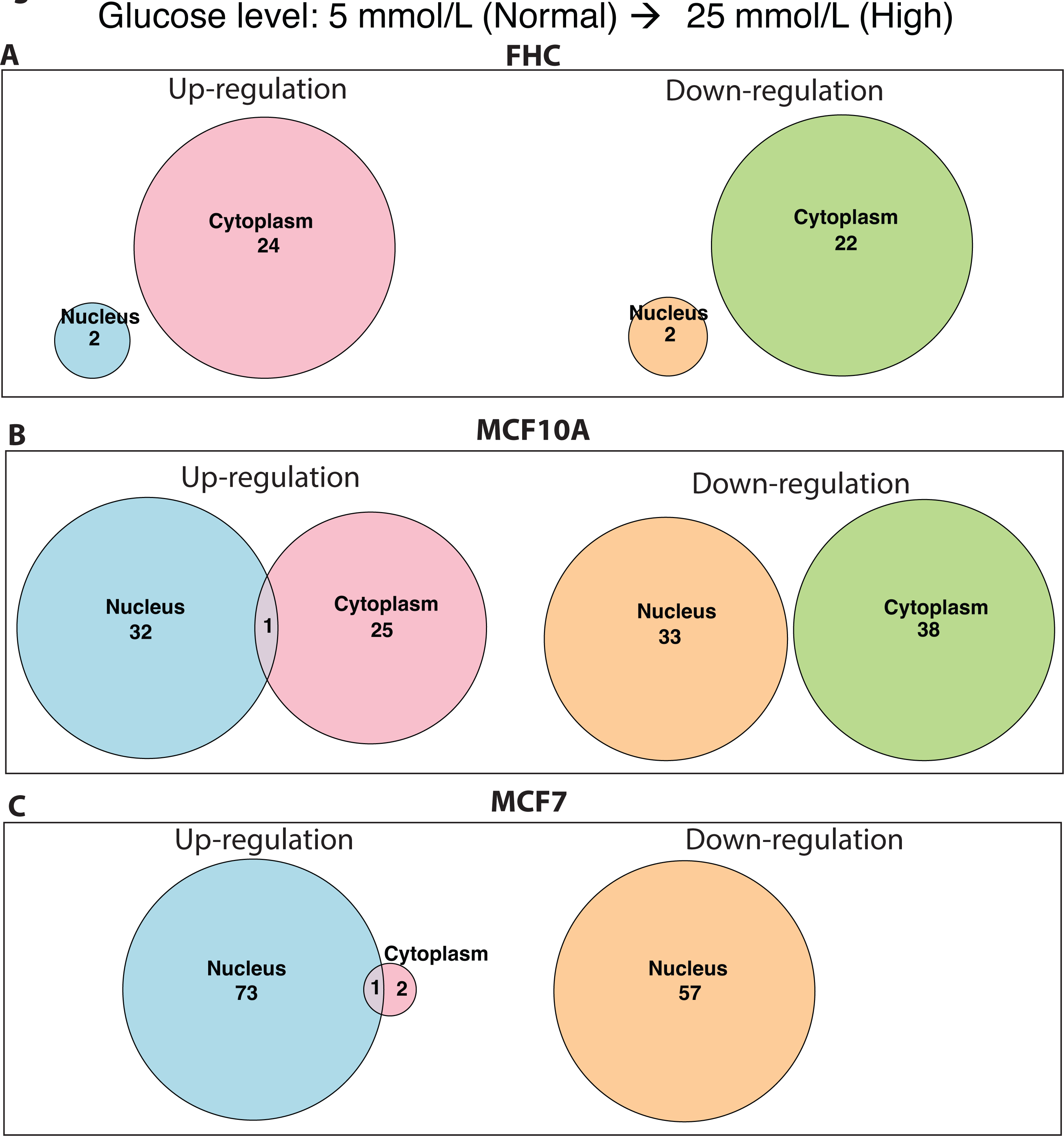
Overlap of upregulated and downregulated genes in response to glucose level changes between the nucleus and cytoplasm. The number of genes upregulated or downregulated in response to an increase in the glucose level from 5 mmol/L to 25 mmol/L for FHC (A), MCF10A (B), and MCF7. (C) cells is indicated in each Venn diagram. The number shown in overlapping arcs indicates the number of genes whose expression was commonly changed between the nuclear and cytoplasmic compartments.

For instance, in the MCF7 cell line, while 74 genes were upregulated in the nucleus under high glucose compared to normal glucose, only one of these genes were similarly upregulated in the cytoplasm. Likewise, 57 genes were downregulated in the nucleus in response to high glucose, yet none were downregulated in the cytoplasm. In the MCF10A cell line, 33 genes were up- regulated under high glucose compared to normal glucose, with only one gene showing consistent changes in the cytoplasm.

The observed discrepancy may be attributed to various factors influencing cytoplasmic mRNA levels beyond transcription, such as mRNA export, degradation, and stability. Our findings, which were consistent across all three cell lines, indicate that transcription alone does not account for glucose-induced changes in cytoplasmic mRNA abundance, and therefore, may not directly predict changes in protein levels.

### Isoform abundance is unevenly distributed between the nucleus and cytoplasm

Alternative splicing occurs in the nucleus, where a single pre-mRNA can give rise to multiple isoforms through RNA processing. If all isoforms have an equal chance of being exported from the nucleus to the cytoplasm, the relative abundance of each isoform would remain consistent between these two cellular compartments. In this study, we investigated whether the relative abundance of different isoforms is maintained between the nucleus and the cytoplasm. We utilized RNA-seq mapped reads to exon-exon junctions to estimate the relative abundance of isoforms and detect potential isoform abundance switches. Specifically, we compared isoforms containing a particular exon with those that skip it, employing a hierarchical framework to account for both estimation uncertainty within individual replicates and variability across replicates to detect alternative splicing changes between two conditions (Shen et al. 2014). To quantify the relative abundance of isoforms containing a specific exon, we utilized PSI (Percent Spliced In), a metric that measures the proportion of transcripts that include a particular exon compared to those that exclude it. PSI values range from 0% to 100%, where 0% represents complete exon skipping, and 100% signifies full exon inclusion. For each comparison, significant splicing changes are defined as those with a PSI difference (ΔPSI) greater than 0.2 between two conditions and a false discovery rate (FDR) below 0.05. As shown in **Figure 3A**, an example exon exhibits a higher inclusion level in the cytoplasm compared to the nucleus in the FHC cell line. Specifically, this exon, located in the DAZ interacting zinc finger protein 1 like (*DZIP1L*) gene, which encodes a ciliary basal body protein localized to centrioles, has PSI values of 0.39 and 0.65 in two replicates of the nucleus. In contrast, its inclusion is significantly elevated in the cytoplasm, with PSI values of 0.78 and 0.78 across two replicates. These results suggest that the relative isoform abundance differs between the nucleus and cytoplasm. **Figure 3B** shows an example of decreased exon inclusion in the cytoplasm. This exon, which is located in the disco interacting protein 2 homolog C (*DIP2C*) gene whose alteration is associated with neurological disorders (Li et al. 2022; Yang et al. 2022) and cancer (Larsson et al. 2017) shows PSI values of 0.39 and 0.45 in the nucleus but drops to 0.07 and 0.26 in the cytoplasm, indicating that isoforms containing this exon are depleted in the cytoplasm compared to the nucleus.

**Figure 3.**
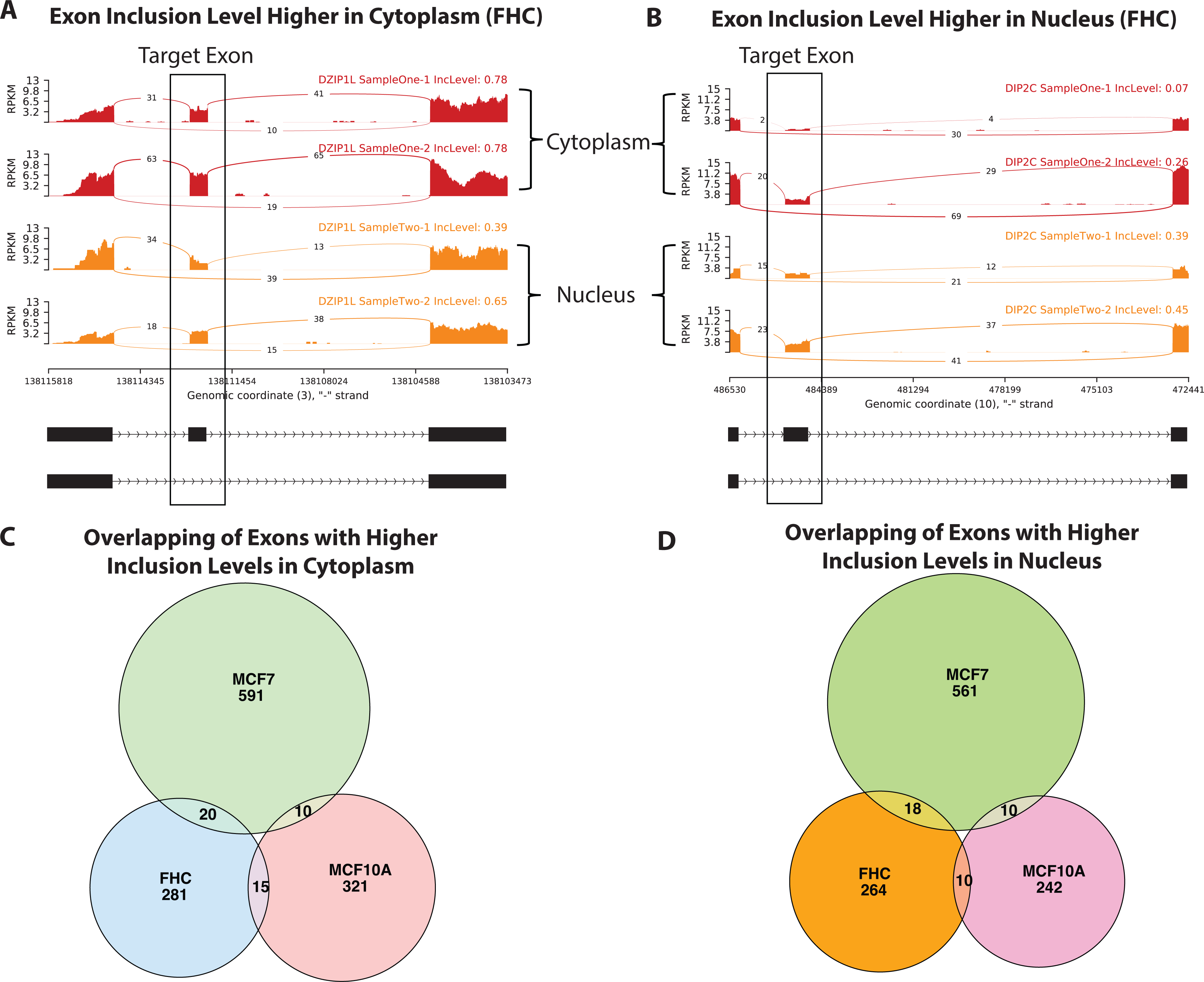
Exon inclusion levels are discordant between the nucleus and cytoplasm. (A) An exon in *DZIP1L* whose inclusion level is higher in the cytoplasm compared with that in the nucleus in FHC cells. (B) An exon in *DIP2C* gene whose inclusion level is lower in the cytoplasm compared with that in the nucleus in FHC cells. (C) Overlapping of exons with higher inclusion levels in the cytoplasm compared with the nucleus among FHC, MCF10A, and MCF7 cell lines. (D) Overlapping of exons with higher inclusion levels in the nucleus compared with the cytoplasm among FHC, MCF10A, and MCF7 cell lines.

In total, we detected 316 and 292 exons with higher inclusion levels in the cytoplasm and nucleus, respectively in FHC cells. To assess whether these isoform preferences in the cytoplasm versus nucleus are cell-type dependent or independent, we conducted a similar analysis in two other cell lines, MCF7 and MCF10A. As shown in **Figures 3C** and **3D**, the isoform preferences appear largely cell-type dependent. For example, among the 316 exons with higher inclusion in the cytoplasm of FHC cells, only 20 exons also exhibit higher inclusion in the cytoplasm of MCF7 cells, and 15 exons in MCF10A cells (**Figure 3C**). Similarly, of the 292 exons with higher inclusion in the nucleus of FHC cells, only 18 exons show higher inclusion in the nucleus of MCF7 cells, and 10 exons in MCF10A cells (**Figure 3D**). These findings indicate that exon inclusion patterns in the cytoplasm and nucleus are not consistent across cell types, highlighting the cell-type specificity of isoform preference between the nucleus and cytoplasm.

### Glucose-induced exon inclusion level changes in the cytoplasm do not mirror those in the nucleus

We further investigated whether glucose-induced exon inclusion level changes observed in the nucleus are consistent with those in the cytoplasm. Across the FHC, MCF7, and MCF10A cell lines, we identified 642, 926, and 643 exons, respectively, that were differentially spliced between normal (5 mmol/L) and high (25 mmol/L) glucose levels in either the nucleus or the cytoplasm. As shown in **Figure 4**, the glucose-induced changes in exon inclusion levels between the nucleus and cytoplasm display minimal correlation. Specifically, the Spearman correlation coefficients (Rho) for the FHC, MCF7, and MCF10A cell lines were -0.176 (P = 0.25), 0.049 (P = 0.69), and 0.228 (P = 0.16), respectively, indicating a lack of significant association.

**Figure 4.**
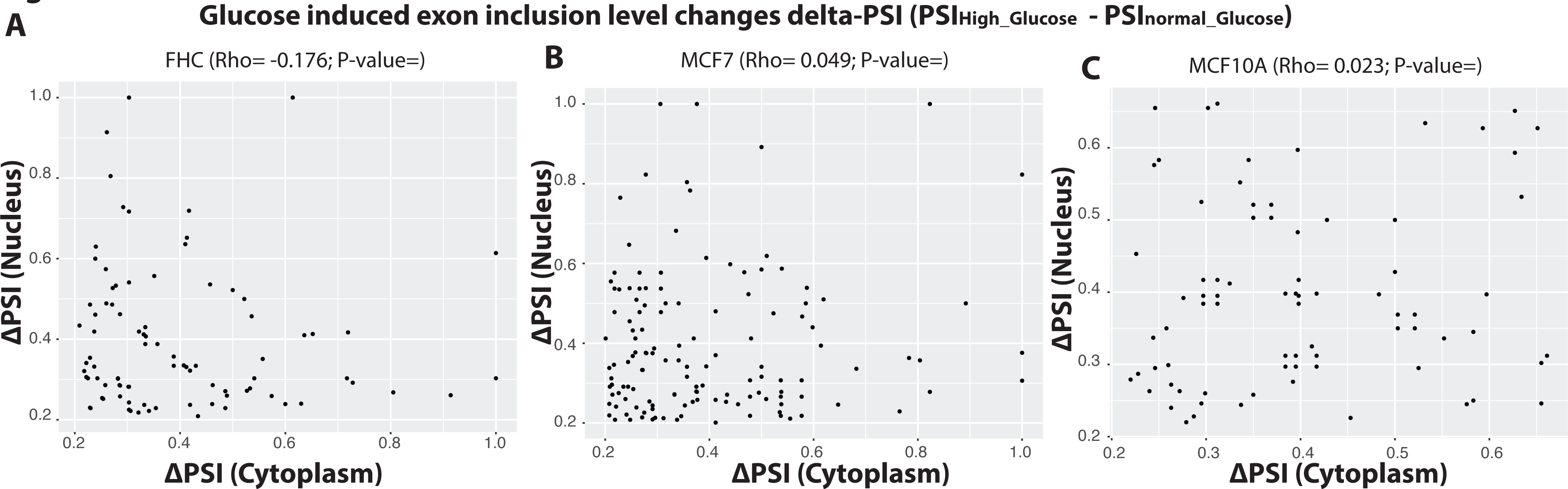
Spearman’s rank correlations of glucose-induced changes in exon inclusion levels between the cytoplasm and nucleus. The Spearman’s correlation coefficient (Rho) is shown for each of the FHC (A), MCF7 (B), and MCF 10A (C) cell lines.

For instance, in the FHC cell line (**Figure 4A**), several exons exhibit substantial inclusion changes in the nucleus (high ΔPSI) but remain largely unchanged in the cytoplasm. A similar trend is observed for MCF7 and MCF10A cells (**Figure 4B, 4C**), where no consistent pattern emerges between the two cellular compartments. These findings suggest that glucose-induced alternative splicing in the nucleus and potential isoform selection during export to the cytoplasm jointly contribute to the complexity of post-transcriptional regulation.

### Pathways Enriched in Glucose-induced gene-level and isoform-level changes are compartment- and cell type-dependent

To determine whether the enriched pathways associated with glucose-induced gene-level and isoform-level changes are dependent on cellular compartment and cell type, we compared enriched pathways on differentially expressed genes (DEGs) and genes with differentially spliced exons (ΔPSI) under normal (5 mmol/L) and high (25 mmol/L) glucose conditions across the FHC, MCF7, and MCF10A cell lines.

As shown in **Figure 5A**, pathways enriched by glucose-induced DEGs are highly depending on the cellular compartment (nucleus vs. cytoplasm) and cell types. For instance, the “HDL remodeling” pathway is only significantly enriched (FDR < 0.05) in the cytoplasm of MCF7 cells, but not in the nucleus or other cell types. Notably, pathways involved in lipid metabolism, immune signaling (e.g., “Interleukin-10 signaling”), and cell cycle regulation showed enrichment patterns that differed between the nucleus and cytoplasm, as well as across the three cell lines.

**Figure 5.**
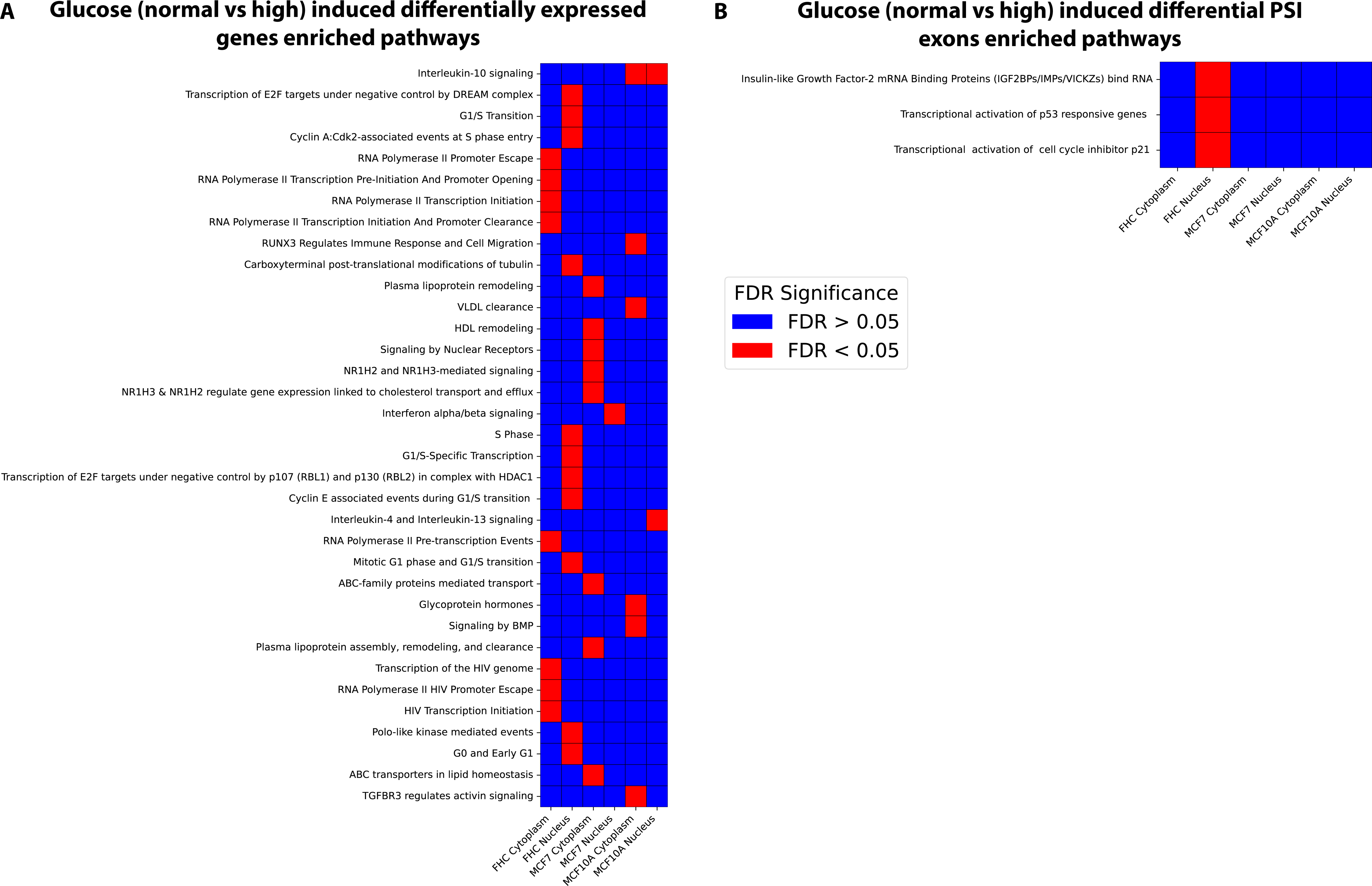
Pathway enrichment analysis of glucose-induced differentially expressed and differentially spliced genes. (A) Pathways enriched in glucose-induced differentially expressed genes. (B) Pathways enriched in genes with glucose-induced differentially spliced exons.

In contrast to DEGs, the pathways enriched by differentially spliced exons (ΔPSI) displayed a more limited but distinct pattern (**Figure 5B**). These splicing changes were highly dependent on both the cellular compartment and the cell line. For example, in FHC cells, genes with alternative splicing under high glucose conditions enriched pathways related to “Transcriptional activation of p53 responsive genes”, “Transcriptional activation of cell cycle inhibitor p21”, and “Insulin-like Growth Factor-2 mRNA Binding Proteins (IGF2BPs)” in the nucleus while the cytoplasm of FHC cells and other cell types did not show such enrichment. This suggests that glucose-induced splicing changes may selectively impact pathways, with unique patterns in different cellular compartments and the cell types.

## Discussions

Most transcriptomic studies rely on bulk RNA sequencing, which measures mRNA levels in whole cells without distinguishing between nuclear and cytoplasmic compartments. This approach assumes that transcriptomic responses are primarily transcriptional. However, our study provides a more nuanced perspective by separately analyzing nuclear and cytoplasmic transcriptomes. We demonstrated that glucose-induced changes in mRNA abundance show minimal concordance between these compartments, highlighting the critical role of post-transcriptional processes, such as mRNA export, stability, and degradation, in shaping the cytoplasmic transcriptome. This discrepancy underscores the limitations of interpreting bulk RNA-seq data as purely reflective of transcriptional responses, as cytoplasmic mRNA levels are shaped by complex post-transcriptional regulatory mechanisms.

In addition to examining mRNA abundance, we explored differences in exon inclusion levels between the nucleus and cytoplasm. Our findings reveal that exon inclusion patterns are highly compartment- and cell-type specific. For example, some exons showed higher inclusion in the cytoplasm compared to the nucleus, while others were preferentially included in the nucleus.

These differences were observed across the three cell lines: FHC, MCF10A, and MCF7, suggesting that alternative splicing and isoform selection during mRNA export contribute significantly to the transcriptomic diversity. Furthermore, glucose-induced changes in exon inclusion often diverged between the two compartments, emphasizing the distinct regulatory landscapes governing nuclear and cytoplasmic transcriptomes.

While our study provides novel insights into the compartment-specific dynamics of glucose-induced transcriptomic changes, it is important to acknowledge its limitations. Our conclusions are based on glucose-induced conditions, and the extent to which these findings generalize to other stimuli or metabolic contexts remains uncertain. Additionally, we focused on three specific cell lines, which may not fully capture the diversity of transcriptomic responses across different cell types or tissues. Future studies could extend this analysis to other metabolic conditions and a wider range of cell types to determine whether the observed compartmental differences are universally applicable.

In summary, our study highlights the critical need for compartment-specific transcriptomic analyses to fully understand the regulatory complexity of glucose-induced cellular responses.

## Materials and methods

### Cell culture and treatment

The human colon epithelial cell line FHC, mammary gland epithelial cell line MCF 10A, and breast cancer cell line MCF7 were purchased from the American Type Culture Collection (ATCC, Manassas, VA). The FHC cells were cultured in DMEM/F-12 medium (Thermo Fisher Scientific, Waltham, MA) supplemented with 10% (v/v) bovine calf serum, additional 10 mM HEPES, 10 ng/ml cholera toxin, 5 μg/ml insulin, 5 μg/ml transferrin, 100 ng/ml hydrocortisone, and 20 ng/ml human recombinant EGF (MilliporeSigma, Burlington, MA). The MCF 10A cells were maintained in DMEM/F-12 medium supplemented with 5% (v/v) horse serum (Thermo Fisher Scientific, Waltham, MA), 100 ng/ml cholera toxin, 10 μg/ml insulin, 0.5 mg/ml hydrocortisone, and 20 ng/ml human recombinant EGF (MilliporeSigma, Burlington, MA). The MCF7 cells were maintained in DMEM medium (Thermo Fisher Scientific, Waltham, MA) supplemented with 10% (v/v) bovine calf serum (MilliporeSigma, Burlington, MA), 1% (v/v) nonessential amino acid and 1% (v/v) sodium pyruvate. All cells were cultured in media containing 100 units/ml penicillin and 100 µg/ml streptomycin (Thermo Fisher Scientific, Waltham, MA) at 37 °C in a humidified incubator with 5% CO2. Cell media were replaced every 2-3 days and subcultured at 75%–80% confluency.

For glucose treatment, cells were plated in a 6-well plate and were allowed to adhere overnight in normal glucose media with 5 mM glucose. The next day, the media were replaced with fresh media containing either 5 mM glucose (low/normal glucose) or 25 mM glucose (high glucose). These concentrations mimic physiological normal and diabetic levels of glucose and were used in previous *in vitro* studies (Corbett 2018; Dutta et al. 2024; Khawkhiaw et al. 2024). The normal- and high-glucose media were prepared by adding the same set of supplements as the culture media and two varying amounts of glucose to glucose-free basal media. Following the media change, the cells were incubated for 72 hours under normal culture conditions.

### RNA extraction

After the incubation period of 72 hours, the cells were harvested for RNA extraction. Total RNA was isolated with on-column DNase I digestion using the E.Z.N.A.® Total RNA Kit I and the E.Z.N.A.® RNase-Free DNase I Set (Omega Bio-tek, Norcross, GA) according to the manufacturer’s instructions. The nuclear and cytoplasmic RNA extraction with on-column DNA removal was conducted using the Cytoplasmic and Nuclear RNA Purification Kit and the RNase-Free DNase I Kit (Norgen Biotek, Thorold, Canada) following the manufacturer’s instructions.

The RNA extracted was stored at − 80 °C until use. All RNA samples passed quality control for the purpose of RNA sequencing at the sequencing facility (Novogene, Sacramento, CA).

### RNA library preparation and sequencing

Library preparation and sequencing were conducted by Novogene (Sacramento, CA). Briefly, messenger RNA was purified using poly-T oligo-attached magnetic beads, and after fragmentation, the first strand cDNA was synthesized using random hexamer primers, followed by the second strand cDNA synthesis. The remaining overhangs of double-stranded cDNA were converted into blunt ends via exonuclease/polymerase activities. After adenylation of the 3’ end of the DNA fragments, sequencing adaptors were ligated to the cDNA, and the library fragments were purified with the AMPure XP system (Beckman Coulter, Brea, CA). The adaptor-ligated cDNA was then amplified using PCR, followed by purification of the PCR products. The quality of the library generated was assessed on the Agilent Bioanalyzer system, and sequencing was performed on an Illumina NovaSeq 6000 platform (Illumina, San Diego, CA) with 150-base-pair paired-end read lengths and more than 20 million read pairs generated per sample.

### RNA-seq data analysis at gene level

RNA-seq Reads were mapped to the human genes (Ensembl v86) using Bowtie (Langmead et al. 2009) allowing up to 3-mismatches and a maximum of 100 multiple hits. The gene expected read counts and the transcripts per million (TPMs) were estimated by RSEM (Li and Dewey 2011). TPMs were median-by-ratio normalized (Leng et al. 2013), and replicates were merged via calculating average normalized TPMs.

### Differentially expressed genes (DEGs)

The EBSeq package(Leng et al. 2013) was used to assess the probability of gene expression (mRNAs) being differentially expressed between any two given conditions. We required that DEGs should have FDR<5% via EBSeq and >2 fold-change of “normalized read counts+1”.

### Identification of Differentially Spliced Exons

We used rMATS (replicate Multivariate Analysis of Transcript Splicing), a statistical method and tool to identify differential alternative splicing events from replicate RNA-Seq data(Shen et al. 2014). This approach maps RNA-Seq reads to exon-exon junctions to quantify exon inclusion levels. rMATS employs a hierarchical framework to account for both estimation uncertainty within individual replicates and variability across replicates. In this study, we focused exclusively on cassette exons, a common type of alternative splicing event in which an exon can be either included or skipped in the mature transcript. Statistically significant differentially spliced exons were defined based on a false discovery rate (FDR) threshold of <0.05 and a difference in exon inclusion levels exceeding 0.2.

### Availability of data and materials

We have submitted the RNA-seq data and the gene expression data (TPMs and Mapping counts) to GEO (accession number: GSE284924).

## Acknowledgments

We thank the support from the Center for Gene Regulation in Health and Disease (GRHD) at Cleveland State University and Prof. Anton Komar. This work was also supported by DARPA AWD00001593 (PJ) and research funding from Biola University (JK).

